# Improved Thermal Stability of Lactoferrin and Oxidative Stability of Iron(II) Sulfate by Co-encapsulation with Low-Methoxyl Pectin

**DOI:** 10.1101/2025.03.07.642131

**Authors:** Claire E. Noack, Tiantian Lin, Younas Dadmohammadi, Yunan Huang, Alireza Abbaspourrad

## Abstract

Complex coacervates between bovine lactoferrin (LF) and low-methoxyl pectin (LMP) were prepared to encapsulate and deliver Fe(II) as an iron fortification ingredient in food, with the goal of enhancing the protein thermal stability and minimizing Fe(II) oxidation. The impact of pH and biopolymer ratio was investigated in terms of complex yield and iron encapsulation efficiency. The thermal stability of LF at 95 °C was determined by changes in turbidity, particle size, and LF retention by HPLC. The interaction between LF and LMP prevented thermal aggregation and circular dichroism analysis showed the complexation preserved the secondary structure of the protein. LF-LMP-Fe complexes contained 28-74 mg g^−1^ of total iron depending on the pH of formation, with <5% of the Fe released at neutral pH. The complex coacervation system shows promise as a thermally stable ingredient for iron delivery and food fortification.

## 1. Introduction

Iron deficiency is the most common nutrient deficiency, affecting more than one-third of the population worldwide (Han et al., 2022). Iron deficiency is most prevalent in middle and low-income countries, children, menstruating women, and in populations that eat a monotonous, plant-based diet (Zimmermann et al., 2005). The first line of treatment for iron deficiency is usually oral supplementation, however, supplementation of iron can cause gastrointestinal distress (Tolkien et al., 2015). Another option is to fortify food with bioavailable iron sources, however, these can cause undesirable off-flavors and color changes in foods (Blanco-Rojo & Vaquero, 2019).

Incorporation of iron ions into a delivery system can mitigate these issues. Encapsulation prevents the interaction between iron and other components of the food matrix by providing a physical barrier (Hurrell, 2002; Tan et al., 2023). Previous approaches to iron ion encapsulation have used proteins, carbohydrates, or lipid-based ingredients as the iron carriers (Kazemi-Taskooh & Varidi, 2021). Further, the bioavailability of iron can be increased through co-delivery with compounds that improve iron absorption, such as ascorbic acid or bovine lactoferrin (LF) (Huang et al., 2024; Teucher et al., 2004)

LF is an iron-binding glycoprotein in the transferrin family that exhibits anti-bacterial, anti-viral, and anti-cancer activity. One LF molecule can bind two Fe(III iron ions with high affinity (K_D_ ∼ 10^-20^ M) (Baker & Baker, 2004). LF exists in holo-(iron-saturated) and apo (iron-free) states, with native LF having an iron saturation between 15-20% (Wang et al., 2019). Commercially, LF is isolated from bovine milk, where the concentration of LF ranges from 0.03-0.49 g L^-1^ (Cheng et al., 2008). LF has been shown to increase iron absorption in adults and infants, likely due to LF-binding receptors in the duodenum (Cox et al., 1979; Mikulic et al., 2020; Suzuki et al., 2005). In addition to binding Fe(III), LF increases the solubility of free Fe(III) in higher pH conditions, which increases the bioavailability of supplemented iron (Uchida et al., 2006). Iron supplementation using LF results in less gastrointestinal distress and fewer side effects than high-dose ferrous sulfate (Zhao et al., 2022).

The thermal stability of LF depends on both iron saturation and pH. At neutral pH, apo-LF has a denaturation temperature of 70 °C, while holo-LF denatures at 85 °C (Bokkhim et al., 2013; Sreedhara et al., 2010). LF is more stable in acidic conditions, with higher retained iron-binding ability and antimicrobial activity after thermal treatment at pH 4 compared to pH 6 (Abe et al., 1991). Improving the thermal stability of LF would allow it to be used alongside Fe supplementation in a broader array of food applications, including foods that undergo thermal processing at neutral pH.

Electrostatic complex coacervation between oppositely charged biopolymers has been used to prevent aggregation and denaturation after thermal treatment (Turgeon et al., 2007). Complex coacervation is a liquid-liquid phase separation where the overall surface charge of particles in solution approaches zero, and repulsive forces between particles are minimized, facilitating aggregation (Kizilay et al., 2011). Complex coacervation systems with LF have been studied, which have included both proteins and polysaccharides (Lin et al., 2022, 2023; Liu et al., 2018).

Pectin is an anionic heteropolymer composed primarily of galacturonic acid monomers commercially extracted from fruits (Lara-Espinoza et al., 2018). The degree of methylation of the galacturonic acid units determines the charge density, with low-methoxyl pectin (LMP) carrying a higher charge density than high-methoxy pectin. LMP has been shown to form electrostatic complexes with LF, protecting the protein from gastric digestion and improving thermal stability (Bengoechea et al., 2011; Niu et al., 2019). Furthermore, pectin is able to interact with divalent cations, including Fe(II) (Celus et al., 2017). Iron and pectin form an association following the egg-box model, where a divalent cation cross-links the anionic polymer chains (Grant et al., 1973). This cross-linking results in a gel which can be used to improve the absorption of iron by slow release under gastric conditions (Ghibaudo et al., 2018). Pectin, along with other probiotic fiber, is thought to improve iron bioavailability by lowering the pH of the duodenum, as well as slowing the speed of digestion (Husmann et al., 2022). Finally, gel formation with pectin can prevent the oxidation of Fe(II) to Fe(III), improving iron solubility and bioavailability (Kyomugasho et al., 2017; Maire du Poset et al., 2018).

Based on this, we hypothesized that the oxidative stability of iron and the thermal stability of the LF would be improved by co-encapsulating LF and Fe(II) with LMP. Therefore, Fe(II)-loaded complex coacervates between LF and LMP were developed. The encapsulation system was optimized by changing the pH of the solution and the mass ratio of biopolymers. The thermal stability of the LF and oxidative stability of the Fe(II) were measured after heating. The complex with the highest amount of LF, and most thermally stable, was then characterized by intrinsic fluorescence, Fourier-transform infra-red spectroscopy and circular dichroism. The optimized complex showed improved thermal and oxidative stability that would enable it to be used as an iron source in food.

## 2. Methods

### 2.1 Materials

Native bovine lactoferrin (Bioferrin 2000; Iron 15 mg 100 g^−1^) was purchased from Glanbia Nationals, Inc. (Fitchburg, WI, USA). Low-methoxyl pectin (LMP) (non-amidated, 35% degree of esterification (DE), extracted from citrus peel) was obtained from CP Kelco (San Diego, CA, USA). Ammonium acetate (>97%)was purchased from Research Products International (Mt. Prospect, IL, USA). Ferrozine (>98%) was purchased from Cayman Chemical Company (Ann Arbor, MI, USA). L-ascorbic acid (99%), sodium phosphate monobasic monohydrate (>98%), sodium phosphate dibasic heptahydrate (>98%), and ferrous sulfate heptahydrate (>99%) were purchased from Sigma-Aldrich (St. Louis, MO, USA)., HPLC grade acetonitrile (99.9%), trifluoroacetic acid (HPLC grade, 99.5%), sodium hydroxide (>97%), hydrochloric acid (38%), and the Pierce Rapid Gold BCA Protein Assay Kit were purchased from Fisher (Hampton, NH, USA). Milli-Q water (18.2 MΩ cm^−1^) purified with the Millipore system (Sigma, MA, USA) was used for all solutions.

### 2.2 Complex Formation Procedure

The complexes between LF and LMP were formed using the pH first method, where the biopolymer solutions are adjusted to the target pH before mixing. LMP (0.2 w/v%) was hydrated in MilliQ water at 60 °C for 20 min, then allowed to sit overnight at 4 °C. LF (0.2 w/v%) LF was hydrated in MilliQ water at room temperature (∼25 °C) for 2 h. The solutions of LF and LMP were adjusted to the target pH of 4, 5, or 6 using 1 M HCl or 1 M NaOH. Then the solutions were mixed in the volume ratios of 8:1, 4:1, 2:1, or 1:1 LF:LMP. To form the ternary (iron-containing) complexes, 80 μL of 0.5 M FeSO_4_⋅7H_2_0 was added to the combined solutions of LF and LMP at all volume ratios. The solutions were mixed for 1 h at room temperature, then centrifuged at 10,000*g* for 10 min at 20 °C. The pellet was collected, frozen at –20 °C overnight, and freeze-dried (Labcono, Kansas, MO, USA) for 48 h at a moisture collector temperature of – 50 C and a vacuum pressure of –0.049 mBar. The yield of the dried complexes was determined by Eq. **1**:

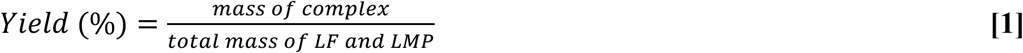

### 2.3 Zeta Potential Measurement

The zeta potential was determined using a Malvern Zetasizer Nano ZS (Malvern, Germany). The LF and LMP solutions (0.2 w/v%) were adjusted to the target pH levels between 2 and 10 at 0.5 increments using 1M HCl or NaOH. The complex solutions were measured without dilution at their mixing ratios. Ternary complexes were measured after the addition of 80 μL of 500 mM FeSO_4_. The measurements were taken using Smoulchwski mode. Each measurement was taken in triplicate with >10 runs per sample. The scale of electrostatic interaction (SEI) was calculated using Eq. **2**:

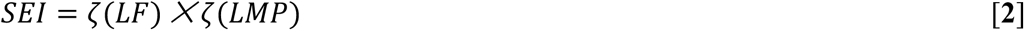

where ζ(LF) and ζ(LMP) are the absolute values of the zeta potential at each respective pH.

### 2.4 Turbidity

Turbidity was determined by measuring the %T of an undiluted sample at 600 nm on a Shimadzu UV/Vis spectrophotometer. The turbidity was calculated according to Eq. **3**:

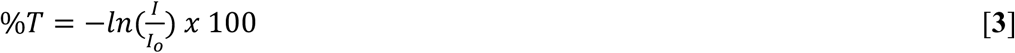

where *I* is the transmittance of the sample and *I_o_* is the transmittance of the blank. MilliQ water (100 %T) was used as the blank.

### 2.5 Particle Size

Particle size was measured at 0.2 w/v% of the LF and LF-LMP-Fe complexes using a Zetasizer (Malvern Nano-Zs90, U.K.). The particle size was determined by dynamic light scattering (DLS) in a disposable 1 cm path length cuvette with a backscattering angle of 173°. The refractive index was set as 1.33 for water and 1.45 for protein-containing samples. Z-average sizes were calculated using cumulants analysis. The analysis was performed in triplicate with at least 11 runs for each sample.

### 2.6 LF Encapsulation and Loading

The LF concentration of the supernatant after complex formation was measured using the bicinchoninic acid (BCA) method, with bovine serum albumin (BSA) as the standard. The encapsulation efficiency was calculated by Eq. **4**:

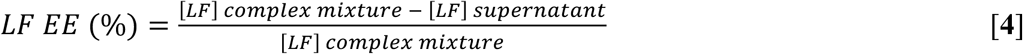

LF loading was determined by High Pressure Liquid chromatography (HPLC) with a reverse phase BioZen Intact XB-C8 HPLC column (150×4.6 mm, 3.6 μm; Phenomenex, Torrence, CA, USA) using the method described by (Lin et al., 2023). The protein complex (1 mg mL^-1^) was hydrated in a pH 7 10 mM phosphate buffer and analyzed at 214 nm using a diode array detector. The LF loading in the sample was determined by Eq. **5**:

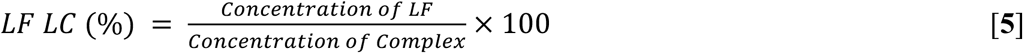

### 2.7 Iron Encapsulation/Loading

Iron encapsulation was determined by using the FerroZine method (Carpenter & Ward, 2017) to measure the concentration of iron remaining in the supernatant after complex formation colorimetrically. An FeSO_4_ standard curve was measured from 0.02 mM to 0.1 mM and used to calculate the Fe concentration of the samples. The Fe encapsulation efficiency (EE) Eq. **6** and loading capacity (LC) Eq. **7** were calculated as follows:

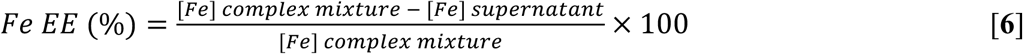

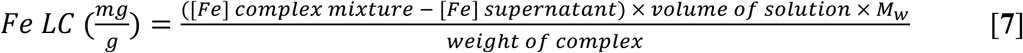

### 2.8 Iron oxidation

The percentage of Fe(II) oxidized to Fe(III) iron was determined by modifying the Ferrozine method. Fe(II) is measured by omitting the ascorbic acid in the first step, and adding 0.2 M HCl. The Fe(II) is compared to the total iron content measured after the addition of ascorbic acid, which reduces Fe(III) to Fe(II). The percent of iron oxidation is calculated by Eq. **8**:

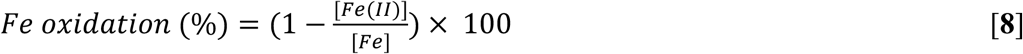

### 2.9 Iron release

The release of iron from the complex was measured after stirring at 500 rpm for 15 min in MilliQ water at 0.1 w/v% and again after heating to 95 °C for 5 min. The iron release percentage was calculated from the concentration of iron measured in the ultrafiltrate using a 3000 kDa molecular weight cut-off (MWCO) centrifugal filter unit via Eq. **9**:

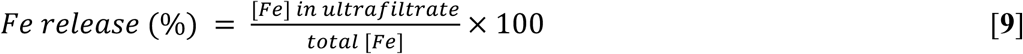

### 2.10 Thermal Treatment and LF retention

LF and LF-LMP complex samples were rehydrated in 10 mM pH 7 sodium phosphate buffer for 15 min prior to loading into glass tubes. The tubes were sealed and the samples heated in a 95 °C water bath for 6 minutes 15 s (the internal temperature of the sample reached 95 °C in 1 min 15 s). The samples were then cooled in an ice bath to room temperature. LF retention was determined by the change of LF concentration before and after heating as quantified by HPLC analysis and calculated using the following Eq. **1****0**:

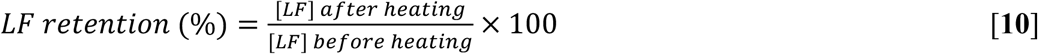

### 2.11 Intrinsic fluorescence spectroscopy

The intrinsic fluorescence of LF, LF and 2 mM FeSO_4_, and LF in binary and ternary complex solutions was measured at pH 4, 5, and 6. The intrinsic fluorescence signal of the protein was measured by excitation at 295 nm and the emission was recorded from 310–400 nm using a Shimadzu RF-6000 spectrophotometer (Shimadzu, Japan) in a 1-cm quartz cuvette.

### 2.12 Circular dichroism spectroscopy

The secondary structure of LF was determined using circular dichroism (CD). 0.002 w/v% samples were analyzed from 260-190 nm in a 1 cm pathlength quartz cuvette using a J-1500 CD Spectrometer (Jasco, Easton MD, USA). The resulting spectra were analyzed using Dichroweb, an online tool for calculating protein secondary structures.

### 2.13 Fourier-transform infrared spectroscopy

The FTIR spectra of freeze-dried LF and LMP, as well as a 1:1 mixture of the two dry biopolymers, and the coacervate formed at pH 6 at a 1:1 ratio of LF to LMP with 2 mM Fe were collected using an IRAffinity-1S Spectrophotometer with a single-reflection attenuated total reflectance (ATR) accessory from Shimadzu (Kyoto, Japan). A background scan was collected before sample collection and an atmospheric correction was performed before sample analysis. The measurement was taken from 500 to 4000 cm^−1^ with an average of 32 scans.

### 2.14 Statistical analysis

All measurements were performed in triplicate. Results are reported as means ± standard deviations. The mean values were compared using the Tukey HSD test, with *p* < 0.05 determining significant differences using JMP Pro16 (SAS Institute, USA).

## 3. Results and Discussion

### 3.1. LF-LMP complex coacervation

The surface charge of each biopolymer changes as a function of the pH of the solution. The zeta potential of 0.2 w/v% LF and LMP solutions were measured from pH 2 to 10 (**Fig 1**) At pH levels above 3, LMP was negatively charged due to the deprotonated carboxyl groups (pK_a_ ∼3.5) of the galacturonic acid units. The LF was positively charged below its isoelectric point (pI) of 8.5. Due to the relatively high pI of LF, there is a large range of pH conditions where it may interact and form complexes with anionic polysaccharides. In the range of pH 3.5–8.5, the positively charged regions on the surface of LF interact with the negatively charged carboxyl groups of the LMP. The SEI increased from pH 2 to 5 due to the increasing negative charge of LMP and decreased above pH 5 due to the diminishing positive charge of LF. This condition is similar to the SEI measured between LF and sodium alginate (SA), another anionic polysaccharide (Wang et al., 2017). Therefore, the optimum range for complex formation was identified as pH 4-6 for further investigation.

**Figure 1.**
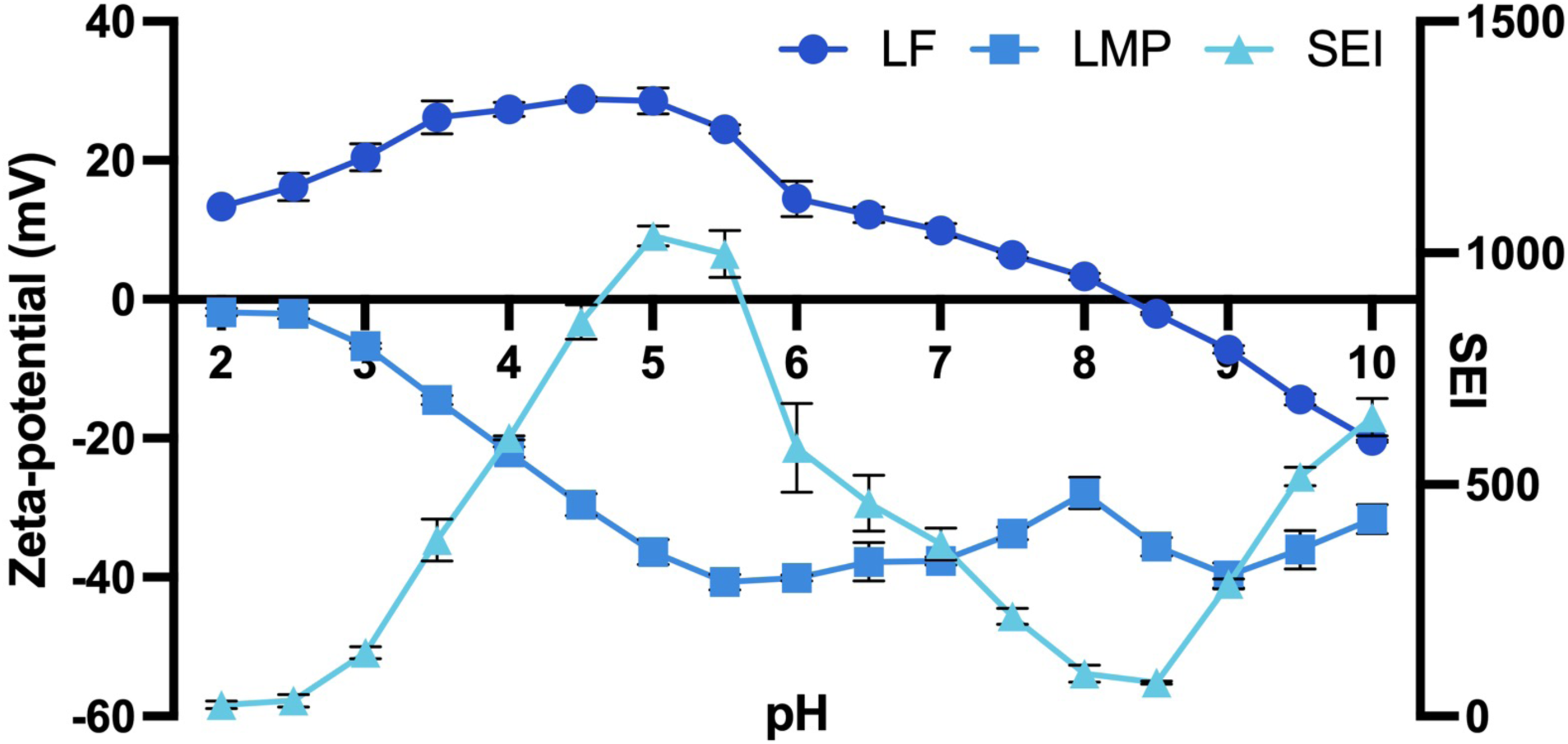
Zeta potential of 0.2 % LF and LMP solutions in MilliQ water. Secondary y-axis shows the SEI at each measured pH.

The effect of different ratios of LF:LMP was investigated at pH 4, 5, and 6. Turbidity, yield, and zeta potential were used to identify the best conditions for complex formation with and without the addition of 2 mM FeSO_4_ **(Fig 2**, **Fig 3)**. LF and LMP are both clear solutions in the range of 4-6, showing no evidence of self-aggregation (**Fig. 2A**). The pH of the mixing solution had a large effect on the turbidity, with pH 4 showing the highest turbidity regardless of the mixing ratio. At pH 5 the turbidity change was smaller, indicating that less complex was formed, which was also confirmed by the decreased yield (**Fig. 2B**). At pH 6 there was almost no increase in turbidity and the solution remained clear, even though the biopolymers are oppositely charged (**Fig 2A**, **Fig 2C**).

**Figure 2.**
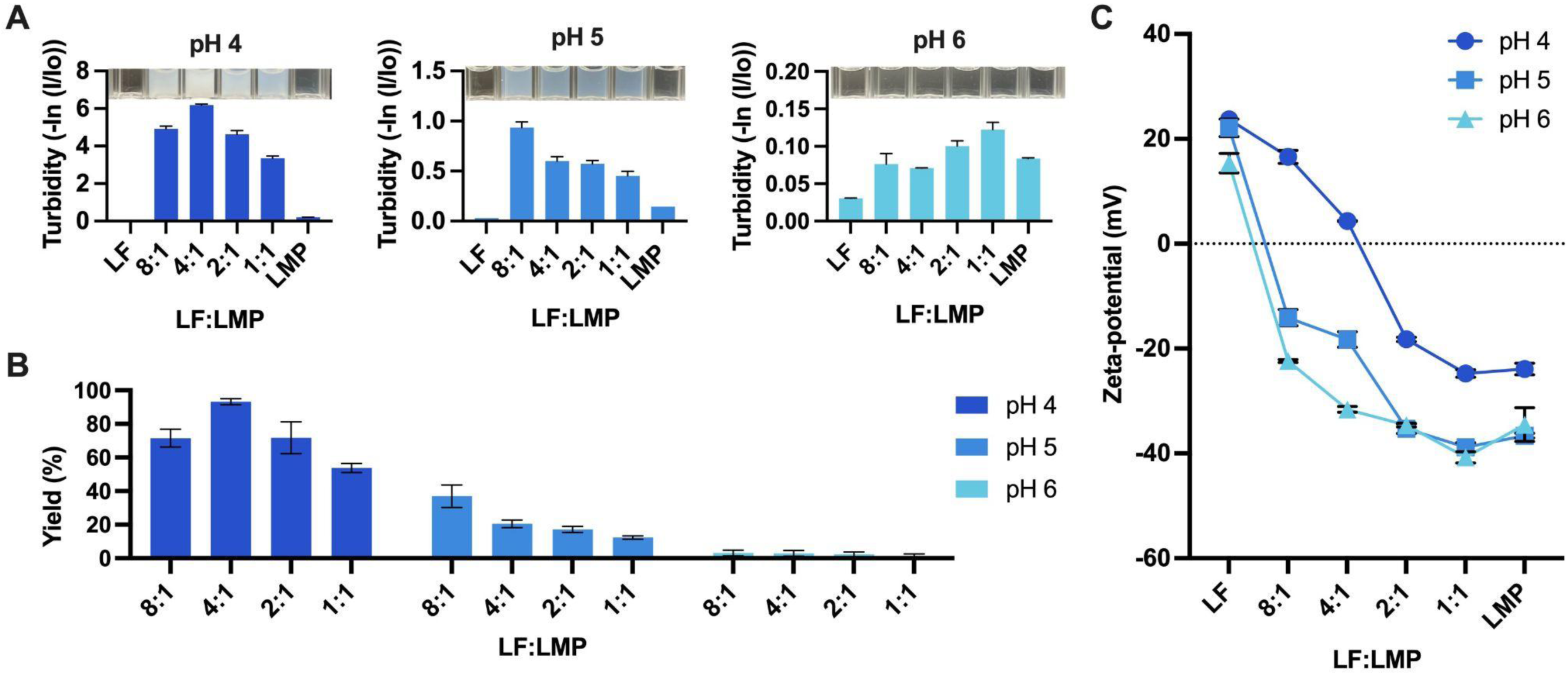
Turbidity of 0.2% total concentration LF, LMP, or LF-LMP mixtures at 8:1-1:1 mass ratios (A). Yield of freeze-dried LF-LMP complexes at pH 4, 5, and 6 (B). Zeta potential of 0.2% LF-LMP mixtures at pH 4, 5, and 6 at different mass ratios (C).

**Figure 3.**
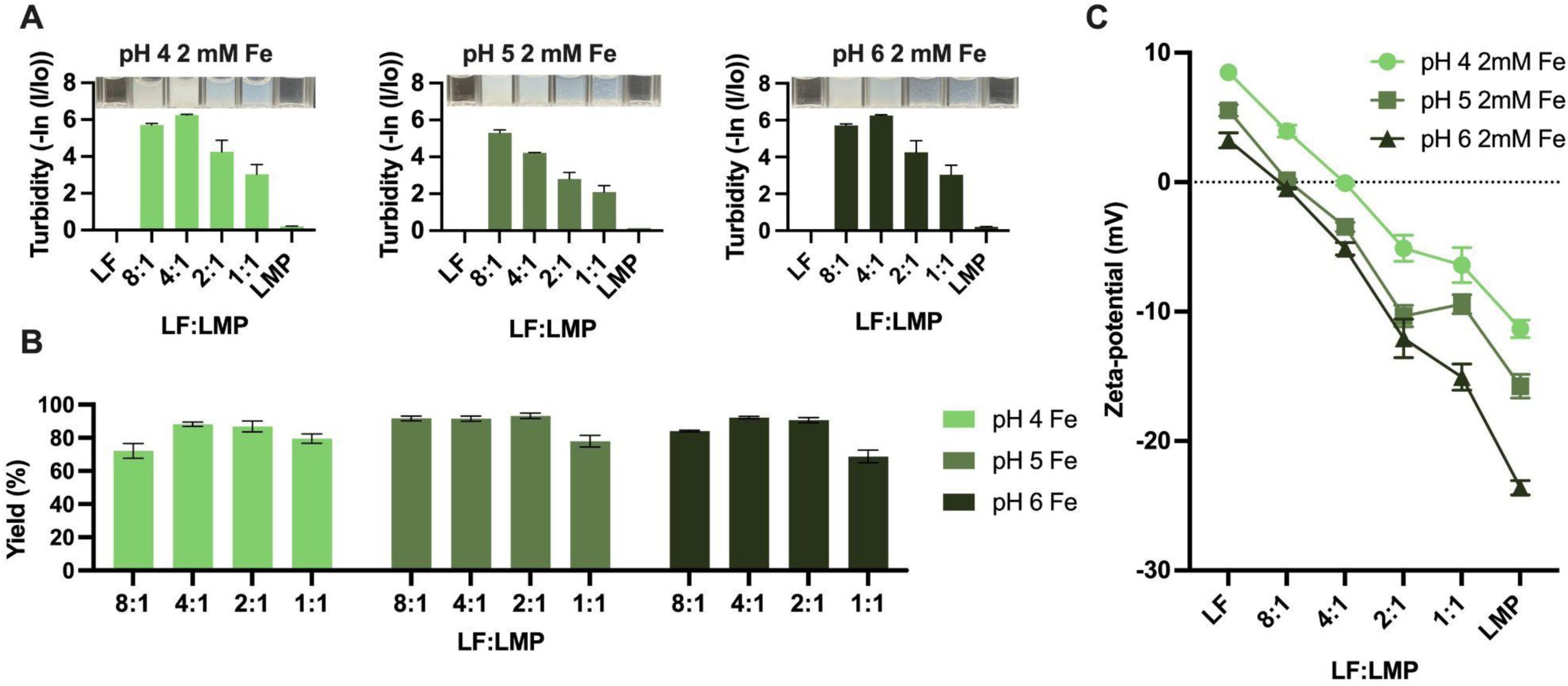
Turbidity of 0.2% total concentration LF, LMP, or LF-LMP-Fe mixtures at 8:1-1:1 mass ratios (A). Yield of freeze-dried LF-LMP-Fe complexes at pH 4, 5, and 6 (B). Zeta potential of 0.2% LF-LMP-Fe mixtures at pH 4, 5, and 6 at different mass ratios (C).

Within each pH, the mixing ratio of LF to LMP impacted the turbidity. At pH 4 the turbidity peaked at the 4:1 mixing ratio, while for pH 5 the highest turbidity was at 8:1. At these mixing ratios, the zeta potential was closest to zero, indicating that the surface charges on both biopolymers were neutralized, forming an insoluble complex. The yield of the complex after centrifuging, collecting the pellet, washing the pellet and then freeze-drying it correlated with the turbidity of the solution mixture (**Fig 2B**). Almost no complex was formed between LF and LMP at pH 6 regardless of the mixing ratio.

When iron sulfate was added the turbidity and yield increased overall (**Fig 3A**, **Fig 3B**). The effect was most dramatic at pH 6, with the yield increasing to 92% at the 4:1 mixing ratio of LF to LMP. The yield of complexes at pH 5 also increased for each mixing ratio. The yield of LF-LMP-Fe complexes at pH 4, however, was lower than LF-LMP at that pH. The effect of the addition of iron to the system on yield can be explained by comparing the zeta potential of the solutions with and without Fe (**Fig 2C**, **Fig 3**). The zeta potential of the 4:1 LF-LMP mixture at pH 4 is nearly zero and is the complex formation condition with the surface charge closest to zero out of any binary complexes formed. The yield of complexes at the other pH and mixing ratios decreases as the surface charge moves further away from zero. The positively charged Fe ions interact with the negatively charged carboxyl groups of the pectin, bringing the measured zeta potential close to zero. The addition of Fe(II) to the system increases the amount of positive charge in the system, driving the surface charge of the particles toward zero. This minimizes the electrostatic repulsion between LF/LMP complex particles, and therefore, the solubility of the complex is decreased.

### 3.2 Bioactives encapsulation and loading

The encapsulation efficiency of both LF and Fe(II) were measured as a function of complex formation pH and biopolymer ratio. The encapsulation efficiency of LF was measured by difference, after quantifying the concentration of LF remaining in the supernatant after the complex was centrifuged and collected by BCA analysis. The encapsulation efficiency of LF correlates with the yield for the binary LF-LMP complexes, reaching nearly 100% for the pH 4 4:1 condition (**Fig 4A**). It is lower for pH 5 LF-LMP and pH 6 LF-LMP, regardless of mixing ratio. The addition of Fe improved LF encapsulation efficiency for pH 5 and 6 complexes, but decreased LF encapsulation for complexes formed at pH 4 (**Fig 4B**). This indicates that at pH 4 LF and Fe compete with each other to interact with the pectin. The increase in LF encapsulation in the ternary complexes compared to binary at pH 5 and 6 is a result of the added positive charge of the iron balancing the more negative charge of the LMP at pH levels below 4.

**Figure 4.**
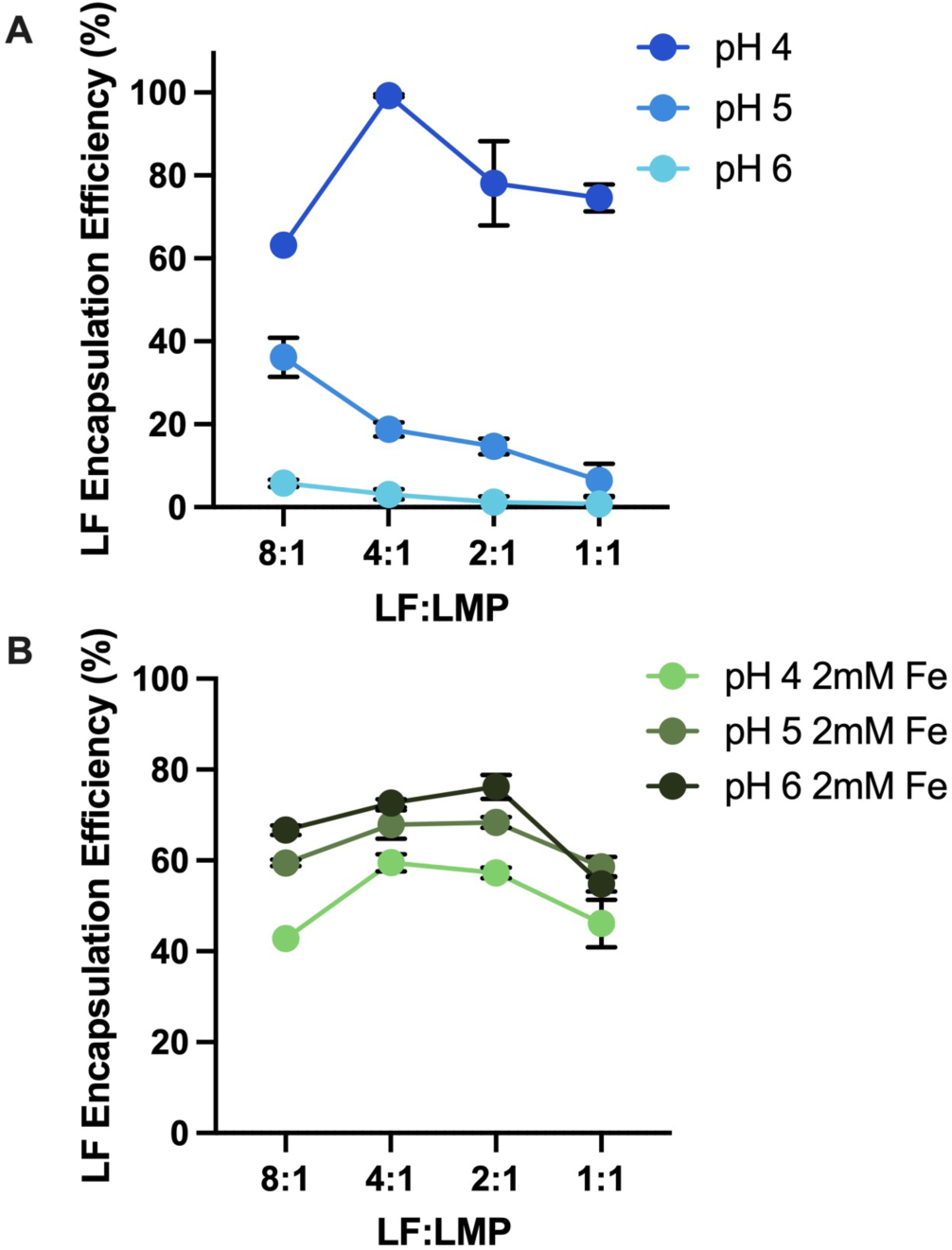
LF encapsulation efficiency of LF-LMP (A) and LF-LMP-Fe (B) complexes calculated by BCA measurement of LF in the supernatant after centrifugation.

The encapsulation of iron is determined by the amount of pectin available. At all pH conditions, the encapsulation efficiency of Fe(II) by the 8:1 mixing ratio of LF to LMP is below 10% but increases with increased pectin concentration (**Fig 5A**). The interaction between pectin and Fe(II) also appears to be pH dependent, with the encapsulation efficiency of the 1:1 ratio of LF to LMP at pH 6 being higher (35%,) than the same solution at pH 4 (17%). This is due to the increase in negative charge density of LMP at higher pH. The stronger interaction between iron and pectin was confirmed by measuring the encapsulation efficiency of LMP and 2mM Fe(II). At pH 4 the encapsulation efficiency of Fe(II) by LMP is 1.9%, but at pH 5 and 6 it is 13.0% and 18.5%, respectively (**Fig S1**). Additionally, the interaction between LF and LMP is weaker at higher pH conditions, potentially allowing more negatively charged carboxyl groups of the pectin to be available for binding Fe(II).

**Figure 5.**
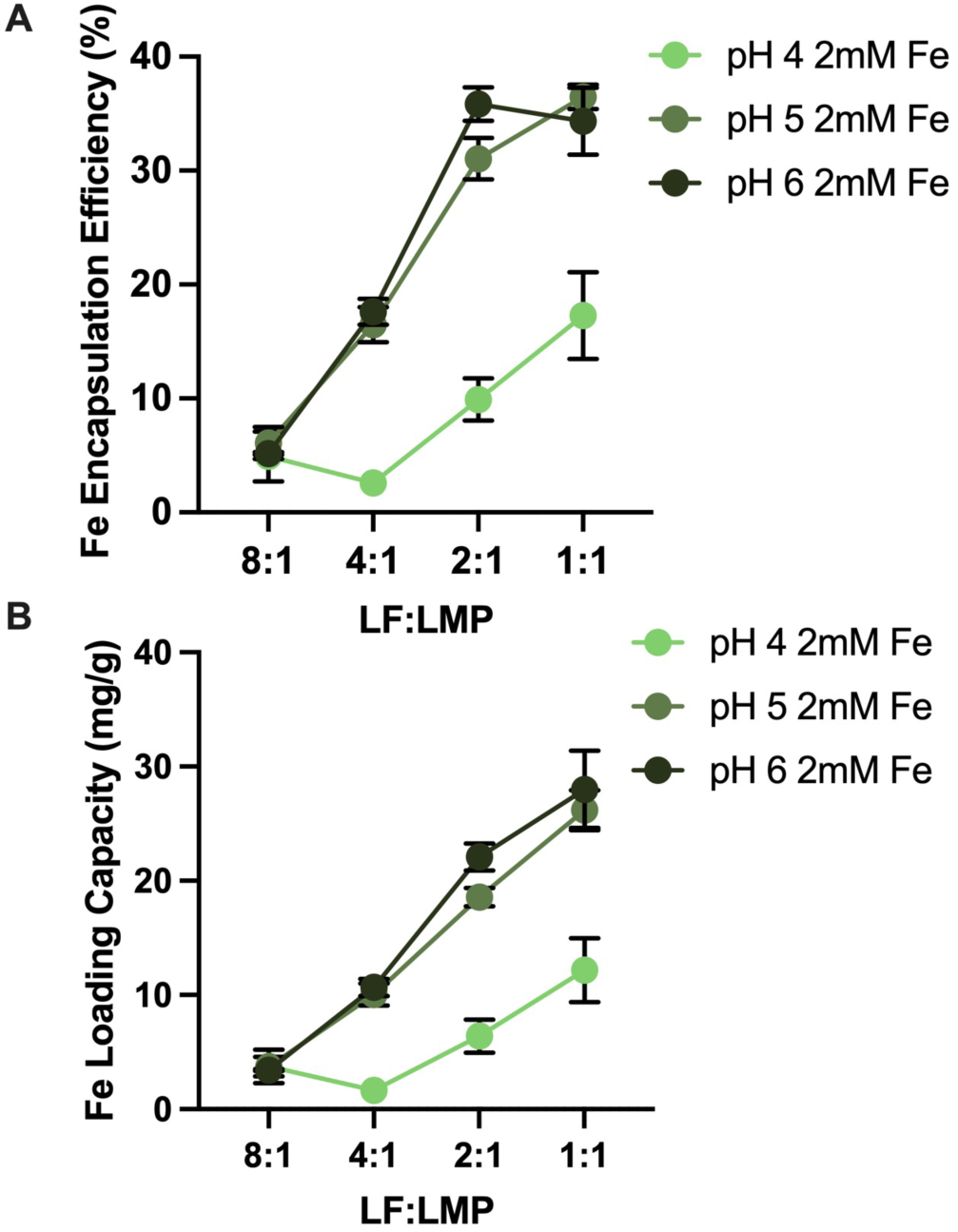
Iron encapsulation efficiency (A) and loading capacity (B) of 8:1-1:1 mass ratio LF-LMP-Fe complexes at pH 4, 5, and 6.

Fe(II) loading capacity depends on both the encapsulation efficiency and yield of the complex. However, due to the similar yield between mixing ratios, it follows the same trend as the encapsulation efficiency. The highest Fe(II) loading occurred with an 1:1 ratio of LF to LMP at pH 5 and pH 6 (**Fig 5A**) where an Fe(II) loading capacity of 26 and 28 mg g^−1^, respectively was observed. This is higher than the Fe(II) loading achieved (13 mg g^−1^) in spray-dried microparticles using whey peptides and maltodextrin (Filiponi et al., 2019). Higher loading has been achieved, however, using a calcium-sodium alginate bead to entrap iron(II) gluconate or ammonium iron(III) citrate (Perez-Moral et al., 2013).

### 3.3 Oxidative stability and release of Fe

Fe(II) is more bioavailable than Fe(III) when used as an oral supplement due to its higher solubility at neutral pH and ability to be transported by the transport protein DMT-1 (Pasricha et al., 2021; Piskin et al., 2022). This makes the preservation of Fe(II) an important goal in iron encapsulation systems. The ratio of oxidized iron to total iron was determined after complex formation (**Fig 6**). The oxidation of iron during the hour-long complex formation is very low, at around 5% for all mixing conditions. Oxidation of iron in the freeze-dried complexes tracks with the pH complex formation, with higher formation pH resulting in higher oxidation. After rehydration in pH 7 MilliQ water, oxidation increases, especially for the pH 6 complex. In comparison to free Fe(II), however, the percent of oxidation in the complex suspensions is significantly lower (p<0.05). After heating at 95 ℃ for 5 min in a solution at pH 7, only half of the Fe(II) in the complex samples was oxidized, but under the same conditions free Fe(II) was 100% oxidized to Fe(III). Therefore, encapsulation of iron within the LF-LMP coacervate inhibits the oxidation of Fe(II) to Fe(III) at neutral pH and under harsh heating conditions, showing proof of concept that a more bioavailable form of Fe can be delivered.

**Figure 6.**
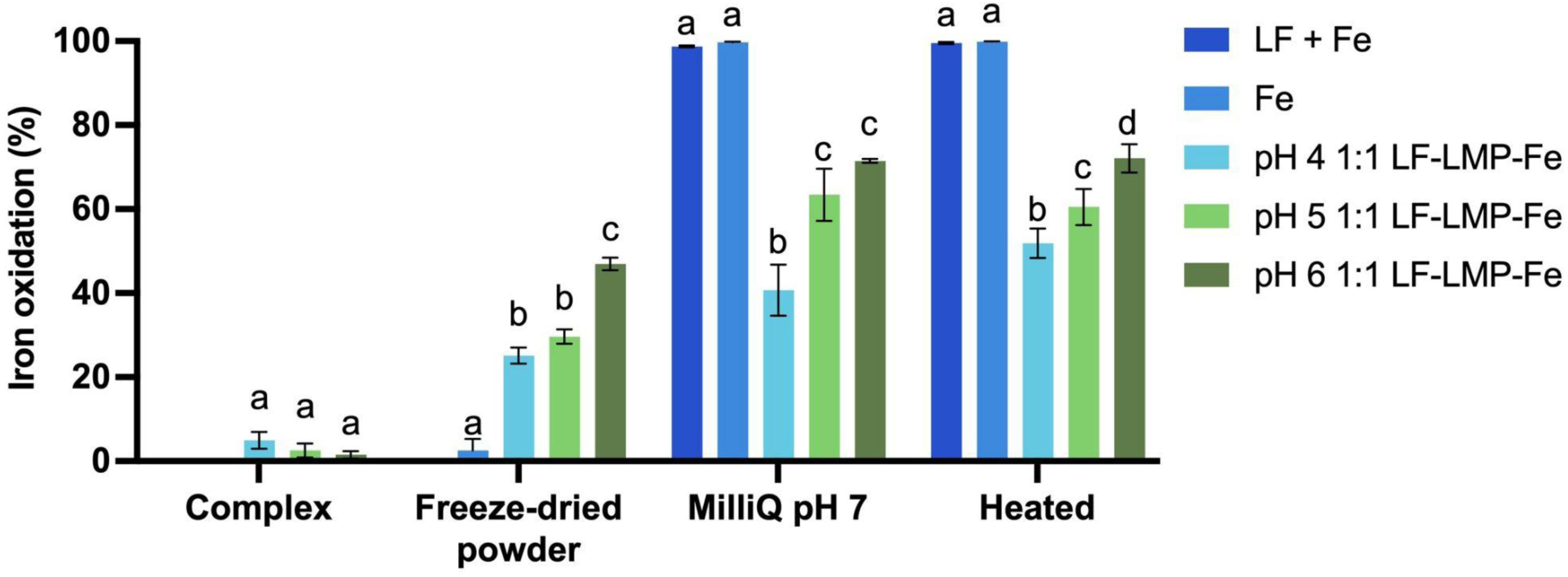
Iron oxidation of 0.5 mM FeSO_4_, 1 mg mL^-1^ LF + 0.5 mM FeSO_4_, or 1 mg/mL LF-LMP-Fe complex prepared at pH 4, 5, or 6 after 1 hour of mixing, after freeze drying, after adjusting to pH 7, and after heating to 95 °C for 5 minutes

The release of iron from the complex was measured after 15 min of hydration time in MilliQ water and again after thermal treatment. The release was measured in an acidic (pH 3) and close to neutral (pH 6) condition. Compared to free Fe(II), significantly less iron (p < 0.05) was detected in the ultrafiltrate from the complexes of a 1:1 ratio of LF to LMP with 2 mM Fe(II) at both pH 5 and pH 6 (**Fig S2**). The thermal treatment had no influence on iron release. However, the iron release was higher in acidic solution compared to neutral solution. This can be explained by the protonation of the pectin carboxyl groups when the solution pH is below the pKa of pectin. With fewer negatively charged carboxyl groups, the Fe(II) was more likely to be released into solution.

### 3.4 Thermal stability of LF

The thermal stability of LF was first measured by changes in the z-average particle size and turbidity before and after heating to 95 °C for 5 min. LF is unstable at neutral pH and denatures between 70 °C and 90 °C, depending on its iron-saturation status (Bokkhim et al., 2013). When heated, LF forms large aggregates, increasing in size from 100 nm to 5 µm, and the turbidity of the solution increases accordingly (**Fig 7A**, **Fig 7B**). These changes in turbidity and increase in particle size are indicative of the thermal denaturation of the LF and the formation of insoluble protein aggregates. In contrast, when redispersed at neutral pH, the freeze-dried complexes did not increase in average particle size or turbidity, remaining dispersed and the solution remained clear. LMP appears to prevent changes in turbidity and particle size even at a low mass ratio within the complex (**Fig S3, Fig S4**). Previous studies have shown that the addition of LMP to LF prevents the formation of aggregates (Bengoechea et al., 2011; Jones et al., 2010).

**Figure 7.**
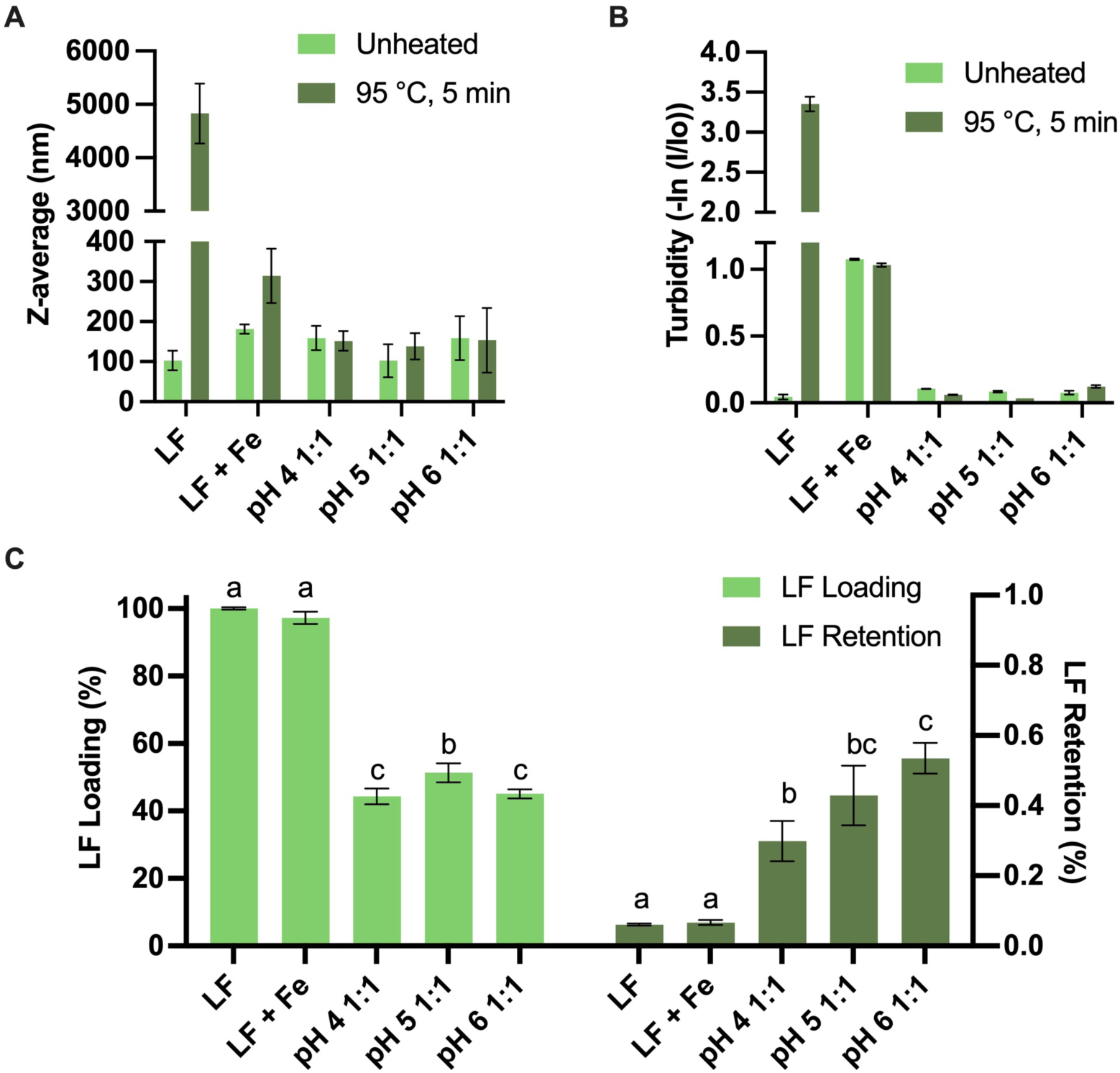
Z-average particle size (A) and turbidity (B) of 0.1% w/v LF-LMP-Fe complexes in 10 mM phosphate buffer before and after thermal treatment. LF loading of LF-LMP-Fe complexes (% w/w) and retention of intact LF after thermal treatment as measured by HPLC (C).

The effect of heating on LF was also measured quantitatively by HPLC. HPLC analysis provided additional information about the complexes that was not noticeable by particle size and turbidity alone. While no ternary complex showed a large increase in particle size or aggregation, both the pH of complex formation and LF loading of the complex had an effect on retention of the LF’s native structure after heating. (**Fig. 7C**). Across all pH complex formation conditions, a higher LF loading resulted in lower native LF structure retention (**Fig. S4**). The retention of the native LF structure with and without 0.5 mM FeSO_4_ was low, 6-7% (**Fig 7**C). Significantly more LF remained in its native structure in the LF-LMP-Fe complexes, more than 30% (p < 0.05). The highest retention (55%) was found for the complex formed at pH 6 and a 1:1 LF to LMP ratio with Fe(II) (**Fig 4C**). Though the LF loading in the pH 6 ternary complex was comparable to the pH 4 complex, the retention is significantly different (p < 0.05). This effect is investigated in the following section, and indicates that the addition of iron to the complex could have a synergistic effect, leading to more stabilizing interactions between the LMP and the LF. The result of the thermal stability study indicated that the optimum protection for LF occurred when the complex was formed at pH 6 with a 1:1 LF to LMP mixing ratio and 2mM FeSO_4_.

### 3.5 Intrinsic Fluorescence of Complexation

To further investigate the interaction between pectin and LF in the presence of iron, the intrinsic fluorescence of the protein was measured (**Fig. 8A-C**). Changes in the intrinsic fluorescence signal reflect changes in protein structure. LF has 13 tryptophan residues that are not solvent exposed and contribute to the intrinsic fluorescence signal of the protein (Schwarcz et al., 2008). The significance of a change in fluorescence intensity is both protein and solvent-specific, and must be experimentally determined by measuring the change in intensity after a perturbation to the structure (Royer, 2006). In proteins with strong internal quenching of the tryptophan fluorescence, a disruption of the structure will cause reduced quenching and an increase in fluorescence. This is the case with LF, where disruption of the native LF structure results in a higher peak intensity (Stănciuc et al., 2013). The addition of Fe(II) to an LF solution caused a decrease in peak intensity, indicating more internal quenching of the tryptophan fluorescence and a more closed structure of the protein (**Fig S6**).

**Figure 8.**
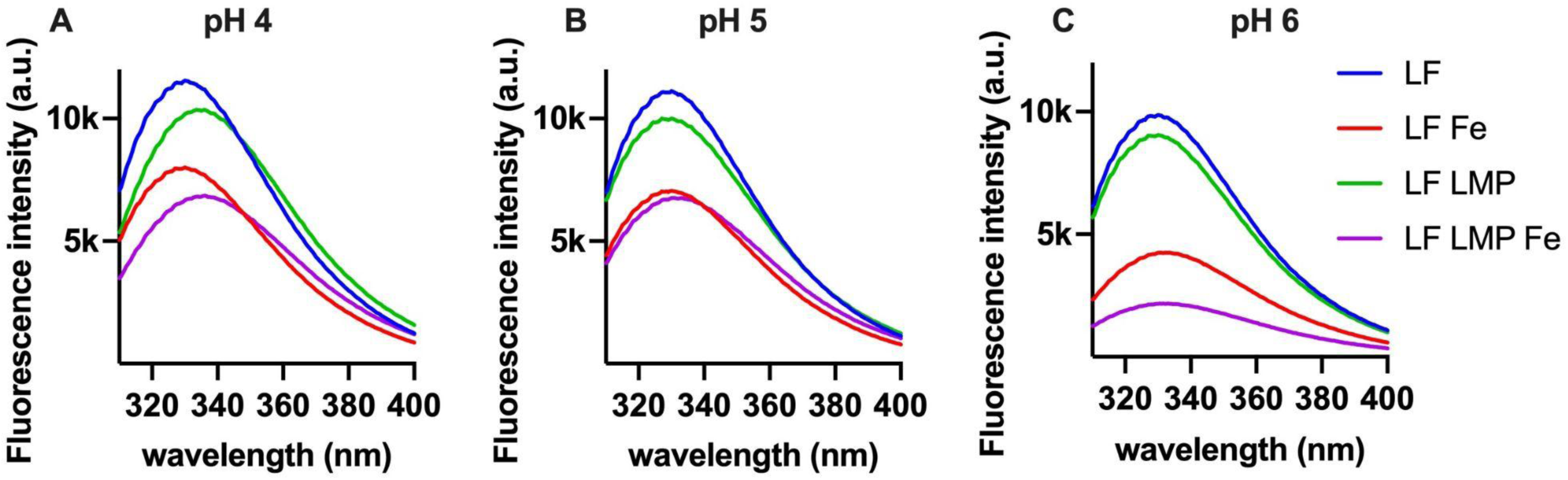
Intrinsic fluorescence of LF (0.1% w/v) at pH 4, 5, or 6 with 0.1% LMP or 2 mM FeSO_4_ in MilliQ water after 1 hour of mixing to form electrostatic complex.

**Figure 9.**
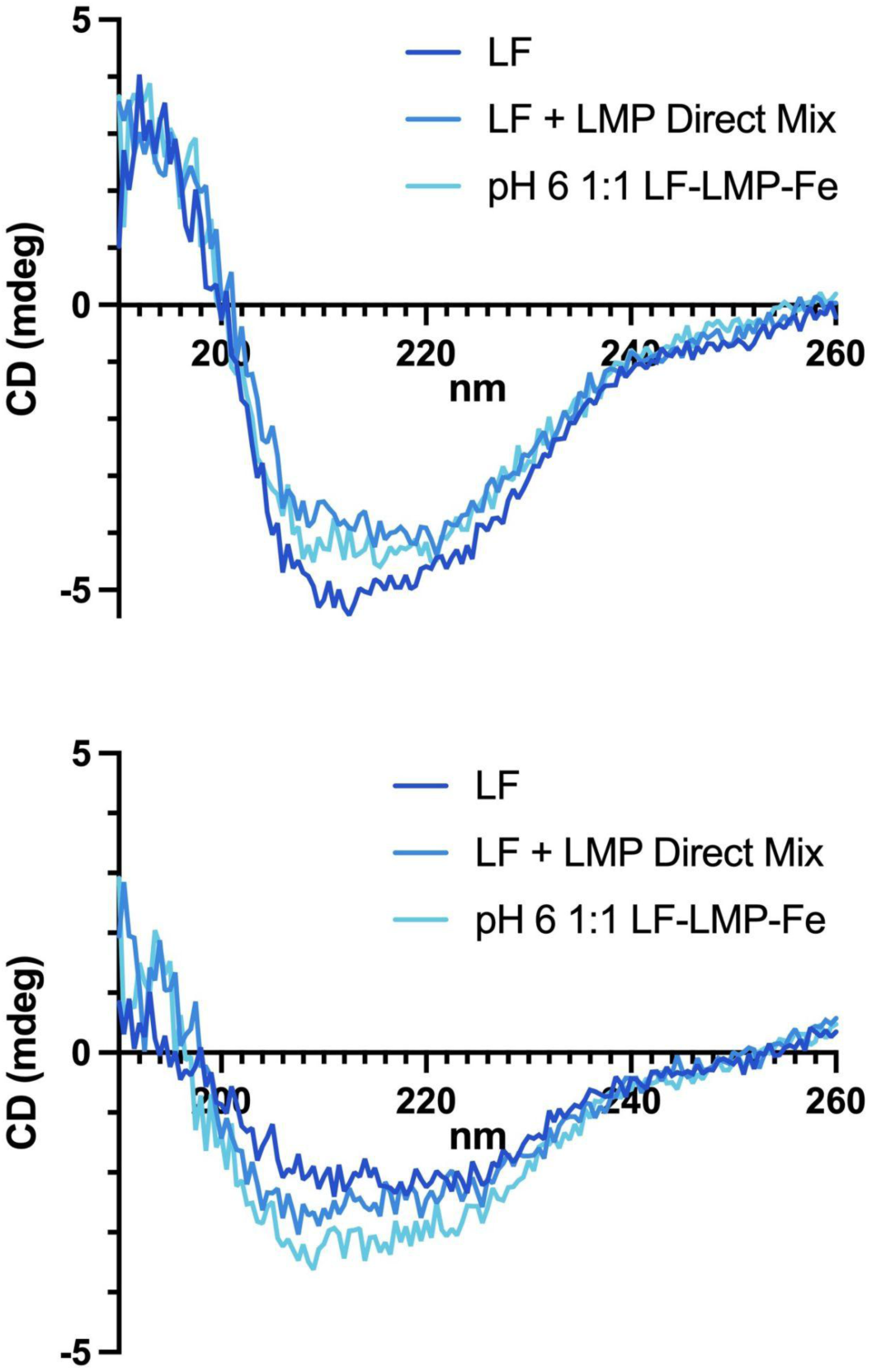
CD spectra of 0.002% LF, LF + LMP + Fe direct mixture, and LF-LMP-Fe complex in 10 mM pH 7 sodium phosphate buffer before (A) and after heating (B).

The intrinsic fluorescence of LF is largely unaffected by pH in the range of 4-6. However, the addition of Fe(II) to a solution of LF results in a decrease in maximum intensity, especially at pH 6 (**Fig 8A-C**). The interaction between LF and LMP in the absence of Fe(II) shows a red shift at pH 4, but no change at pH 5 or 6 **(Fig 8C**). When Fe(II) is added to LF and LMP, the red shift is more noticeable for the pH 4 ternary complex. The largest reduction in maximum intensity occurs in the ternary complex at pH 6. This reduction in peak intensity provides evidence that the addition of Fe(II) promotes the interaction between LF and LMP. This stronger interaction between the polymers in the presence of Fe(II) explains the difference in LF retention between the pH 4 and pH 6 LF-LMP-Fe complexes.

### 3.6 Circular dichroism

Due to the chiral nature of protein structures, the absorption of polarized light in the range of 190-260 nm is characteristic of the makeup of alpha helices, beta sheets, turns, and unordered regions in the protein (Greenfield, 2006). Online tools such as Dichroweb can analyze the spectra and calculate the percentage of each type of sub-structure (Miles et al., 2022).

The CD spectrum of LF after the addition of LMP and the complexation process did not significantly change (p > 0.05) (**Table 1**). Complexation with polysaccharides decreased the calculated % ɑ-helix from 48% to 43%, in line with previous reports that complex coacervation does not disrupt the secondary structure of LF (Lin et al., 2022). Thermal treatment, however, disrupted the secondary structure of LF, resulting in an increase in β-sheet and unordered regions of the protein, and decreasing the percentage of ɑ-helices from 48% to 14%. This is consistent with other reported measurements of LF secondary structure after thermal treatment (Fan et al., 2019). The addition of LMP to the solution prevented some changes in the secondary structure, with 20% ɑ-helix remaining after thermal treatment. The LF was best protected in the LF-LMP-Fe complex, with no significant change in the percentage of any secondary structure element. The retention of the secondary structure indicates the protein was not denatured in the LF-LMP-Fe coacervates during thermal treatment.

**Table 1.** Secondary structure elements of LF and LF complexes in pH 7 10 mM sodium phosphate buffer calculated by DichroWeb before and after thermal treatment.

| Sample | Helix (%) | Strand (%) | Turns (%) | Unordered (%) |
| --- | --- | --- | --- | --- |
| LF | 48 ± 2 <sup>a</sup> | 18 ± 6 <sup>a</sup> | 12 ± 2 <sup>a</sup> | 22 ± 7 <sup>a</sup> |
| LF H | 14 ± 6 <sup>b</sup> | 30 ± 6 <sup>b</sup> | 13 ± 1 <sup>a</sup> | 42 ± 1 <sup>c</sup> |
| Direct Mix | 40 ± 4 <sup>a</sup> | 16 ± 2 <sup>a</sup> | 13 ± 1 <sup>a</sup> | 31 ± 3 <sup>ab</sup> |
| Direct Mix H | 20 ± 1 <sup>b</sup> | 25 ± 1 <sup>ab</sup> | 14 ± 1 <sup>a</sup> | 41 ± 1 <sup>bc</sup> |
| LF-LMP-Fe | 43 ± 2 <sup>a</sup> | 19 ± 5 <sup>ab</sup> | 11 ± 2 <sup>a</sup> | 27 ± 4 <sup>a</sup> |
| LF-LMP-Fe H | 40 ± 2 <sup>a</sup> | 19 ± 1 <sup>ab</sup> | 11 ± 1 <sup>a</sup> | 29 ± 2 <sup>a</sup> |

\*Different letters denote statistically significant (p<0.05) difference between values

### 3.7 Fourier-transform infrared spectroscopy

The complex formation was investigated for changes to chemical bonding by analyzing the FTIR spectra of the biopolymers (**Fig S7**). The amide bands of the protein are visible, with group 1 (1625-1750 cm^−1^), group II (1475-1575 cm^−1^), and group III (1225-1425 cm^−1^) corresponding with previously published FTIR results of LF (Bastos et al., 2018). The spectrum of LMP shows the absorption due to the ester carbonyl group stretching at 1740 cm^−1^ and the carboxylate group stretching from 1630-100 cm^−1^ (Kyomugasho et al., 2017). The spectrum showing a physical mixture of the biopolymers shows no new peaks or major changes when compared to the spectrum of the ternary LF-LMP-Fe complex, indicating that there are no changes in chemical bonding due to the complexation process.This supports the hypothesis that the complex coacervate formation depends on electrostatic interactions, and does not involve covalent bonds that may change the protein activity.

## 4. Conclusion

The synergistic effect of co-encapsulation of LF and Fe(II) by LMP was demonstrated. The Fe(II) encapsulation efficiency was dependent on both biopolymer ratio and pH of the solution at complex formation. The yield of the complex as well as the LF encapsulation efficiency could be dramatically improved by the addition of Fe(II). The addition of LMP to LF prevented thermal aggregation and HPLC analysis shows at the optimized complex formation condition more than 50% of the LF retained its intact structure after harsh thermal treatment at 95 °C for 5 minutes. The optimized condition for ternary complex formation, based on LF retention and Fe(II) loading, was at pH 6 with 1:1 ratio of LF:LMP. The secondary structure of LF was quantified by CD and showed that it was preserved after heating, with no significant difference in the proportion of alpha helices, compared to a significant decrease in alpha helices after thermal treatment in non-complexed LF. The interaction between LMP and Fe(II) was able to prevent the complete oxidation to the Fe(III) form, preserving a more bioavailable form of iron for nutrient delivery. This delivery system could be used to improve iron uptake compared to iron salts alone when used as an iron fortification method.

## Supporting Materials

Supporting information includes additional figures and raw data that was used to generate the figures in this manuscript can be found at: https://doi.org/10.5281/zenodo.14927656

## CRediT authorship contribution statement

**Claire Noack:** Conceptualization, Investigation, Data curation, Formal analysis, Writing – original draft, Writing – review & editing. **Tiantian Lin:** Conceptualization, Methodology, Investigation, Validation, Writing – review & editing. **Younas Dadmohammadi:** Conceptualization, Supervision, Funding acquisition, Writing – review & editing. **Yunan Huang:** Investigation, Writing – review & editing. **Alireza Abbaspourrad:** Project Administration, Funding acquisition, Resources, Writing – review & editing, Supervision.

## Generative AI Statement

The authors certify that generative AI was not used in preparing this article. Non-generative AI, such as spelling and grammar checkers in Office 365 and Google Docs, and citation management software, was used. All instances when non-generative AI was used were reviewed by the authors and editors.

## Acknowledgements

This project was funded by the Bill & Melinda Gates Foundation (INV-039533). The authors acknowledge the use of facilities and instrumentation supported by NSF through the Cornell University Materials Research Science and Engineering Center DMR-1719875.

