## Supplemental Figures for "Improved Thermal Stability of Lactoferrin and Oxidative Stability of Iron(II) Sulfate by Co-encapsulation with Low-Methoxyl Pectin"

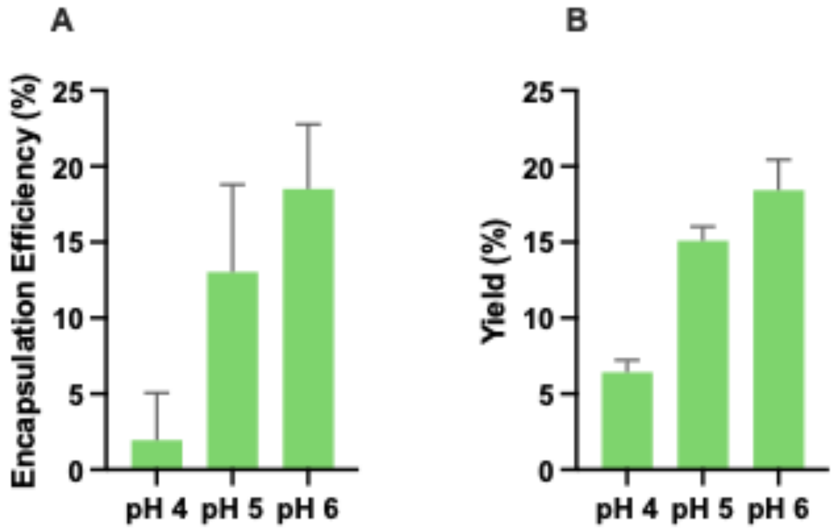


Figure S1. Iron encapsulation efficiency (A) and yield (B) of 0.2% w/v LMP with 2mM $FeSO_4$ in MilliQ water adjusted to pH 4, 5, or 6.

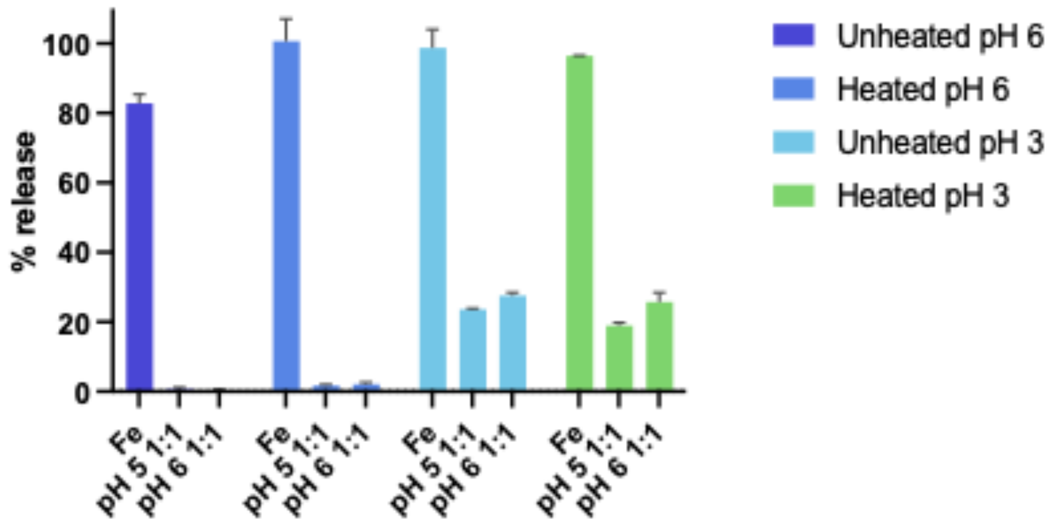


Figure S2. Iron release from pH 5 and pH 6 1;1 LF-LMP-Fe complexes compared to free iron and free iron added to LF

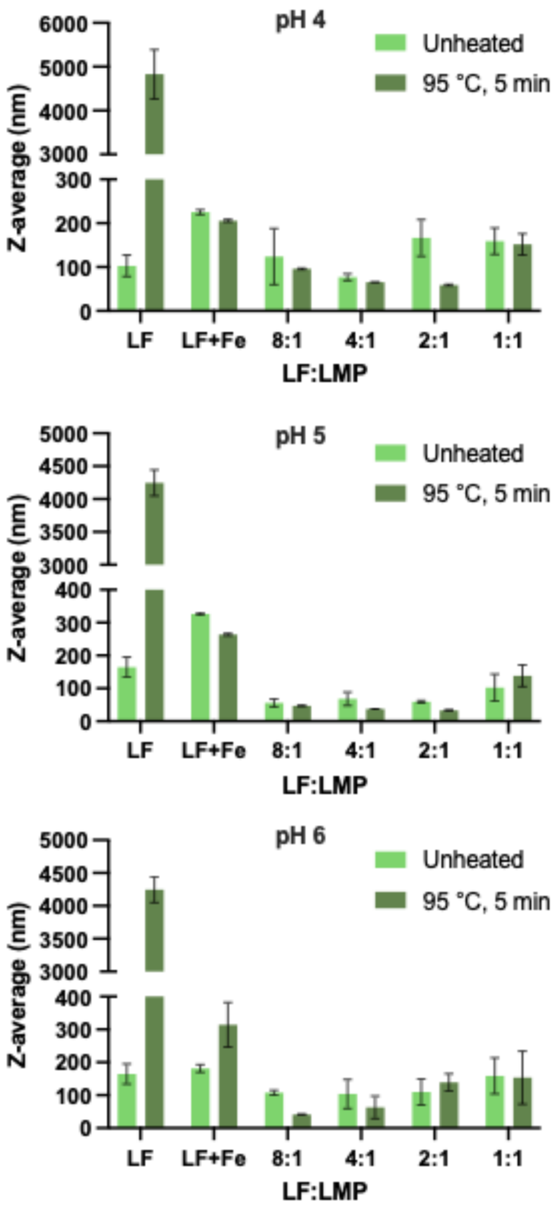


Figure S3. Z-average particle size of redispersed 0.1% w/v 8:1-1;1 LF-LMP-Fe complexes formed at pH 4, 5, or 6 in 10 mM phosphate buffer before and after thermal treatment.

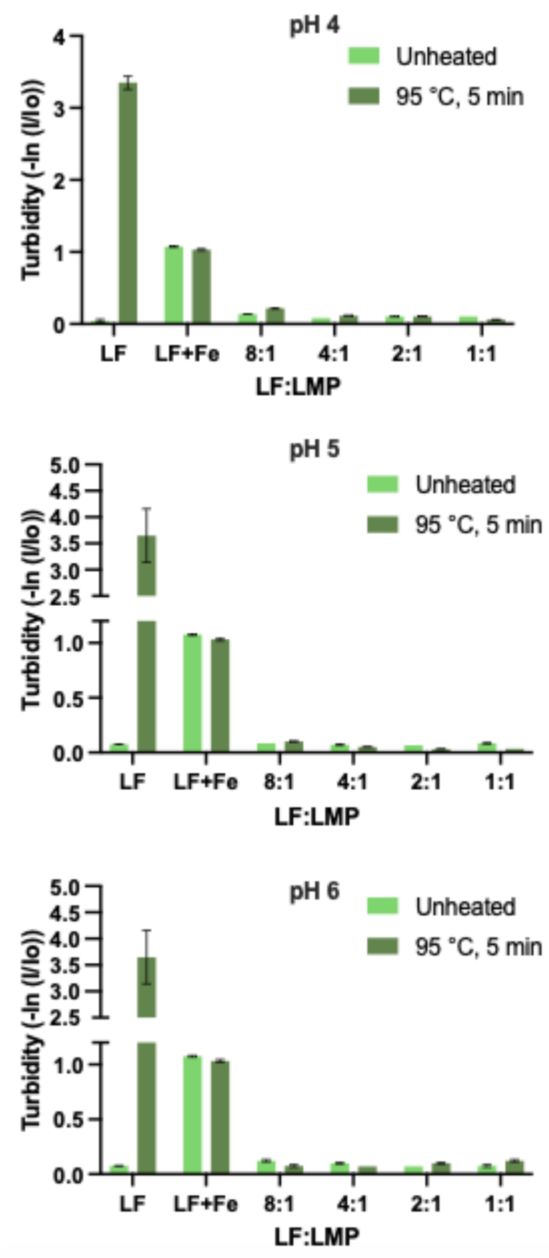


Figure S4. Turbidity of redispersed 0.1% w/v 8:1-1;1 LF-LMP-Fe complexes formed at pH 4, 5, or 6 in 10 mM phosphate buffer before and after thermal treatment.

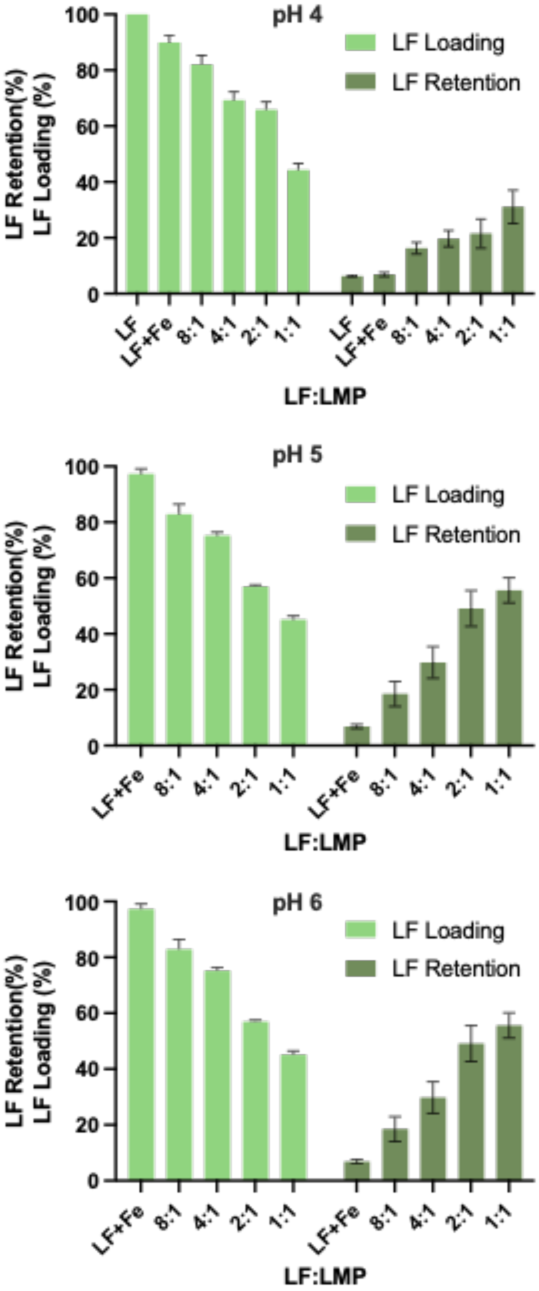


Figure S5. LF loading ratio and LF retention after thermal treatment in pH 4, 5, or 6 LF-LMP-Fe complexes.

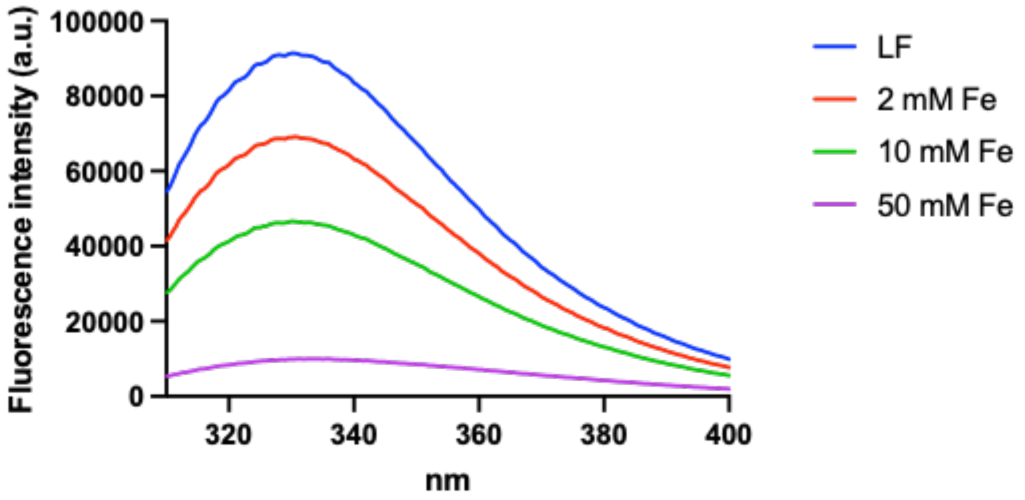


Figure S6. Intrinsic fluorescence of 0.1% LF in MilliQ water with 2-50 mM $FeSO_4$.

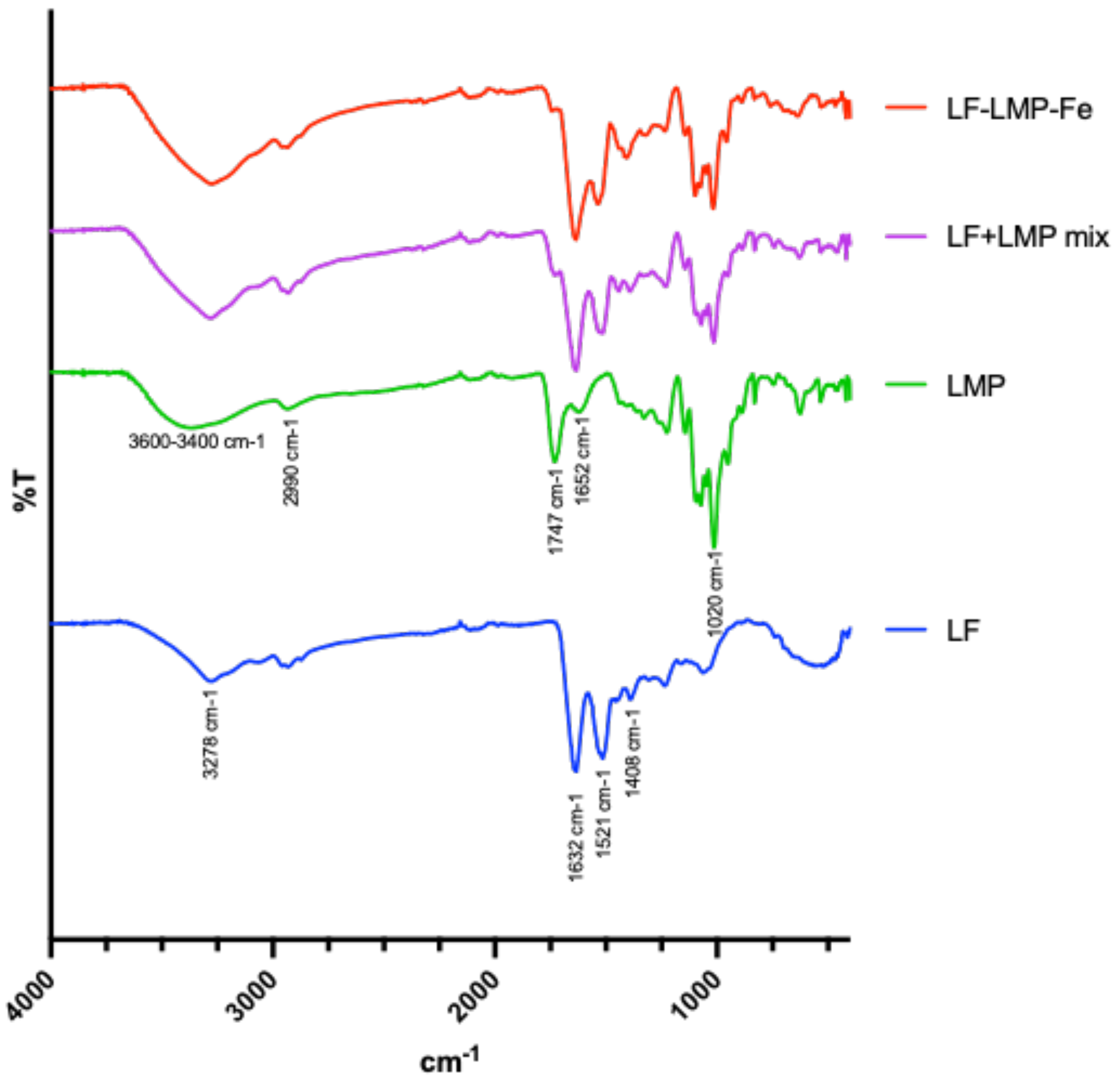


Figure S7. FTIR of LF, LMP. freeze-dried LF-LMP-Fe complex, or a physical mixture of LF and LMP powders.
